# Evaluation of the Physicochemical Characteristics of Honey Produced by Honey Bees *(Apis mellifera)* in Waghimra Zone

**DOI:** 10.1101/2025.03.04.641522

**Authors:** Yesuf Ibrahim, Moges Kibret

## Abstract

Honey is a natural sweet substance produced by bees from plant nectar or secretions. This study focused on assessing the physicochemical properties of honey from the Waghimra Administrative zone in Ethiopia’s Amhara region, and examining how different agro-ecological zones (lowland, midland, and highland) and sampling sources (beekeepers, SDARC apiary site, and local honey traders) influence honey quality. A total of 27 honey samples were collected across these agro-ecological areas. Physicochemical characteristics were analyzed following international standards, including the European Union Directive 2001/110/EC and guidelines from the Ethiopian Standards Agency (2011). The findings showed that the average moisture content, pH, ash content, HMF (hydroxymethylfurfural), free acidity, electrical conductivity, total solids, reducing sugars, and apparent sucrose levels were 17.16%, 3.56, 0.1g/100g, 1.99mg/kg, 24.17meq/kg, 0.25mS/cm, 82.84%, 66.68g/100g, and 4.29g/100g, respectively. All these values were within the acceptable ranges set by the Quality and Standards Authority of Ethiopia, Codex Alimentarius, and European Union standards, confirming the honey’s high quality.

## 1. INTRODUCTION

According to the Codex Alimentarius Commission (2001), honey is defined as a naturally sweet substance produced by bees from the nectar of flowers or the secretions of living parts of plants. The bees collect, transform, combine with their own specific substances, store, and mature this substance in their honeycombs.

Naturally, honey is a supersaturated sugar solution, comprising approximately 95% carbohydrates by dry weight, with fructose and glucose being the main sugar components (Gela et al., 2023). The composition of honey varies depending on the source of the plant, the type of bee, geographical origin, climatic conditions, season, harvesting, processing, and storage conditions (Santos et al., 2023). The exact chemical composition and physical properties of natural honey depend mainly on the type of plants that the bees feed on, climatic conditions, and vegetation, which are important factors that can affect the various properties of honey (Buba, 2013). Most of the volatile compounds in honey include alcohols, ketones, aldehydes, acids and esters, which are also responsible for the taste and aroma characteristics of honey (Maniy-Loh et al., 2011).

Physicochemical properties such as color, moisture, ash, conductivity, pH, free acids, hydroxymethylfurfural (HMF), sugars, sucrose and maltose provide characterization and classification of honey. They are used as criteria for selecting suitable processing and packaging technologies and application of technology for raw honey. It is also important to better respond to consumer demands and to detect and prevent possible manipulation and tampering of honey. It also allows beekeepers in particular and the country in general to receive better prices (Gizaw et al., 2020).

As an important component of the local economy, honey production contributes to the livelihoods of many communities in the Waghimra administrative zone. Ensuring the quality and safety of locally produced honey is important not only to maintain consumer confidence but also to maintain the economic viability of beekeeping in the area (Alemu et al., 2019). This study is in line with broader efforts to support sustainable beekeeping practices and increase the marketability of Ethiopian honey, thereby contributing to the socio-economic development of the Waghimra administrative zone.

To ensure rigorous and standardized assessment, this study was conducted in accordance with established international standards and guidelines (European Union Directive 2001/110/EC, Ethiopian Bureau of Standards, 2011). These benchmarks provide a comprehensive framework for assessing physicochemical parameters, ensuring consistency and comparability when assessing honey quality. Adherence to these standards is essential to obtain reliable and meaningful results that can contribute to broader honey quality research.

Therefore, the objective of this study was to evaluate the quality of honey produced in Waghimra zone based on physicochemical parameters compared to the standards of Ethiopian, European and international honey standards.

## 2. Materials and methods

### 2.1 Honey sample collection

In this study, three honey sampling sources were considered and nine honey samples were collected from each sampling source, using aseptic techniques, resulting in a total of 27 honey samples of approximately 1 kg per sample. The first sampling source was fresh honey collected directly from modern hives. For this purpose, mature honey of *Apis mellifera* was collected from SDARC apiaries in different agro-ecological areas of the Waghimra administrative zone during the honey collection period. The second sampling source was *Apis mellifera* honey samples purchased from local honey traders in each agroecologically based woredas (Sekota - Midland, Gazgibla - Highland and Ziquala - Lowland). The third sampling source was fresh honey directly collected from traditional hives. For this purpose, mature honey of *Apis mellifera* was collected from various agroecological areas of Waghimra administrative zone during the honey collection period. The collected honey samples were transported to Sekota Dryland Agricultural Research Center for physicochemical analysis.

### 2.2 Determination of Physicochemical Analysis of Honey

#### 2.2.1 Color analysis

Honey color was measured using mm P-Fund Honey Color Grading Tool using the approved USDA color standard (USDA, 1985). The P-fund color results of the honey samples were compared with the glycerol analysis standard as per the Codex Alimentarius Commission standard procedure (Codex Alimentarius Commission, 2001).

#### 2.2.2 Moisture Content

The moisture content of honey samples was measured using an Abbe refractometer that was temperature controlled at 20°C and regularly calibrated with distilled water. The honey samples were homogenized and placed in a water bath until all sugar crystals were dissolved. After homogenization, the surface of the refractometer prism was coated with honey and the refractive index was measured for water after 2 minutes. The refractive index values of the honey samples were determined using a standard table developed for this purpose (Bogdanov, 2009).

#### 2.2.3 pH and free acidity

10 g of honey from each honey sample was dissolved in 75 ml of distilled water in a 250 ml beaker and stirred using a magnetic stirrer. The electrode of the pH meter (EUTECH device) was immersed in the solution and the pH of the honey was recorded. Before measuring the honey samples, the device (pH meter) was calibrated with buffer solutions pH 4, pH 7 and pH 9.1. To measure the free acidity, the solution was additionally titrated to pH 8.30 using 0.1 M sodium hydroxide (NaOH). For accuracy, a 10 mL burette was used and readings were recorded to the nearest 0.2 mL. Free acidity was expressed in milliequivalents or millimoles acid/kg of honey and was equal to ml 0.1 M NaOH x 10. Results were expressed to the nearest tenth according to the procedure of the International Honey Commission (Bogdanov, 2009). Acidity = 10 V, where: V = volume of 0.1 N NaOH in 10 g of honey.

#### 2.2.4 Ash content

Determination of ash content was carried out by incinerating honey samples at 600°C in a muffle furnace to constant mass. First, the ash crucibles were heated in an electrical muffle furnace at ashing temperature and subsequently cooled in a desiccator to room temperature and weighted to g (M2). Then 5 g (M0) of each honey sample was weighed to the nearest 0.001 g and taken into a platinum crucible and two or three drops of olive oil was added to prevent foaming. Water was removed and started ashing without loss at a low heat rise to 350 – 400^°^C using a hot plate. After the preliminary ashing, the crucibles were placed in the preheated furnace and heated for at least 4-6 hrs. The ash crucibles were cooled in the desiccators and weighed. The ashing procedure was continued until a constant weight was reached (M1). Lastly, % of the weight of ash in g/100 g honey was calculated using the following formula:

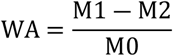

Where M0= Weight of honey taken. M1= Weight of ash + crucible M2= Weight of crucible.

#### 2.2.5 Electrical conductivity

The electrical conductivity of a solution of 20 g dry matter of honey in 100 milliliters of distilled water was measured using an electrical conductivity cell. A 0.745 g of potassium chloride (KCl), was dried at 130°C, dissolved in freshly distilled water in a 100 ml flask, and filled to volume with distilled water. Forty milliliters of the potassium chloride solution was transferred to a beaker and the conductivity cell was connected to the conductivity meter, the cell rinsed thoroughly with potassium chloride solution and immersed the cell in the solution, together with a thermometer and reading of the electrical conductance of the solution in milli siemen after the temperature has equilibrated to 20°C was taken as described in harmonized international commission.

The cell constant K was calculated using the following formula: K=11.691×1/G

Where: K= the cell constant in cm-1, G= the electrical conductance in mS, measured with the conductivity cell. 11.691= the sum of the mean value of the electrical conductivity of freshly distilled water in mS.cm-1 and the electrical conductivity of a 0.1M potassium chloride solution, at 20°C.

#### 2.2.6 Hydroxyl methyl furfural (HMF)

HMF was determined using UV– spectrophotometer. A 5 g of honey sample was weighed in a small beaker and mixed in 25 ml distilled water and transferred into 50 ml volumetric flask. A 0.5 ml carrezz solution I (15 g K_4_Fe (CN) _6_. 3H_2_O /100 ml distilled water) was added and mixed with 0.5 ml carrezz solution II (30 g Zn acetate /100 ml distilled water). The solution was mixed with the honey solution. A droplet of alcohol was added to the solution. The solution was filtered through a filter paper and the filtrate (10 ml) was discarded. A 5 ml filtrate was added to each of the two test tubes and 5 ml distilled water was added to the first test tube (sample solution), while 5 ml sodium bisulfite solution (0.20% of 0.20 g NaHSO_3_/100 ml distilled water) was added into the other test tube (reference). The contents of both test tubes were well mixed by vortex mixer and their absorbance was recorded spectrophotometrically by subtracting the absorbance measured at 284 nm for HMF in the honey sample solution against the absorbance of reference (the same honey solution treated with sodium bisulfite, 0.2%) at 336 nm and the result was calculated and expressed according to international honey commission.

Hydroxyl methyl furfural (HMF)/100 g honey = (A284 - A336) *149.7 * 5*D/W sample; Where A284= absorbance at 284, A336 = absorbance at 336, 149.7= constant, 5= theoretical nominal sample weight and W= mass of honey sample, D=dilution factor where necessary.

#### 2.2.7 Reducing sugar and apparent sucrose determination

This method is a modification of the Lane and Eynon procedure, involving the reduction of Soxhlet’s modification of Fehling’s solutions. The Layne–Enyon technique as explained in AOAC (<u>1990</u>) was used for the estimation of reducing sugar. Briefly, 5 mL of Fehling’s solution A and B will be taken in a <u>250</u> mL Erlenmeyer flask with 7 mL H_2_O and 15 mL of honey. With this solution, 1 mL <u>0.2</u>% methylene blue indicator was added. Thereafter, titration was continued with heating the solution until decolorization of the indicator.

Amount of sucrose was determined using the inversion process. In short, 50 mL of honey was taken in a <u>100</u> mL volumetric flask in which 10 mL dilute HCl was added followed by heating in a water bath, and volume was made up to the mark. Again, the Layne–Enyon procedure was followed for this solution. Amount of sucrose was calculated using the formula of Saxena (2010). % Sucrose= [Total sugar-Total reducing sugar]×0.95

### 2.3 Data management and statistical analysis

The data was entered to the computer using Microsoft excel and statistical package for social science (SPSS) version 26 was used for the analysis of quantitative data. The honey quality data was analyzed by one way ANOVA using SPSS software.

## 3. RESULT AND DISCUSSION

### 3.1 Results

#### 3.1.1 Physicochemical properties of honey

##### A. Color of honey

Among the honey samples analyzed in the current study, about 33.3 % were white, 55.6% were extra light amber, 7.4% were light amber and 3.7% were amber. The honey samples collected from lowland were white, extra light amber and light amber with respective percentage 11.1%, 14.8%, 7.4%. Honey samples collected from midland were 11.1% white and 22.2% extra light amber. The honey samples collected from the highland of the study area were 11.1% white, 18.5% extra light amber, and 3.7% amber (Table 1). The final color designation of the analyzed honey samples was based on USDA Color Standards Designations.

**Table 1:**
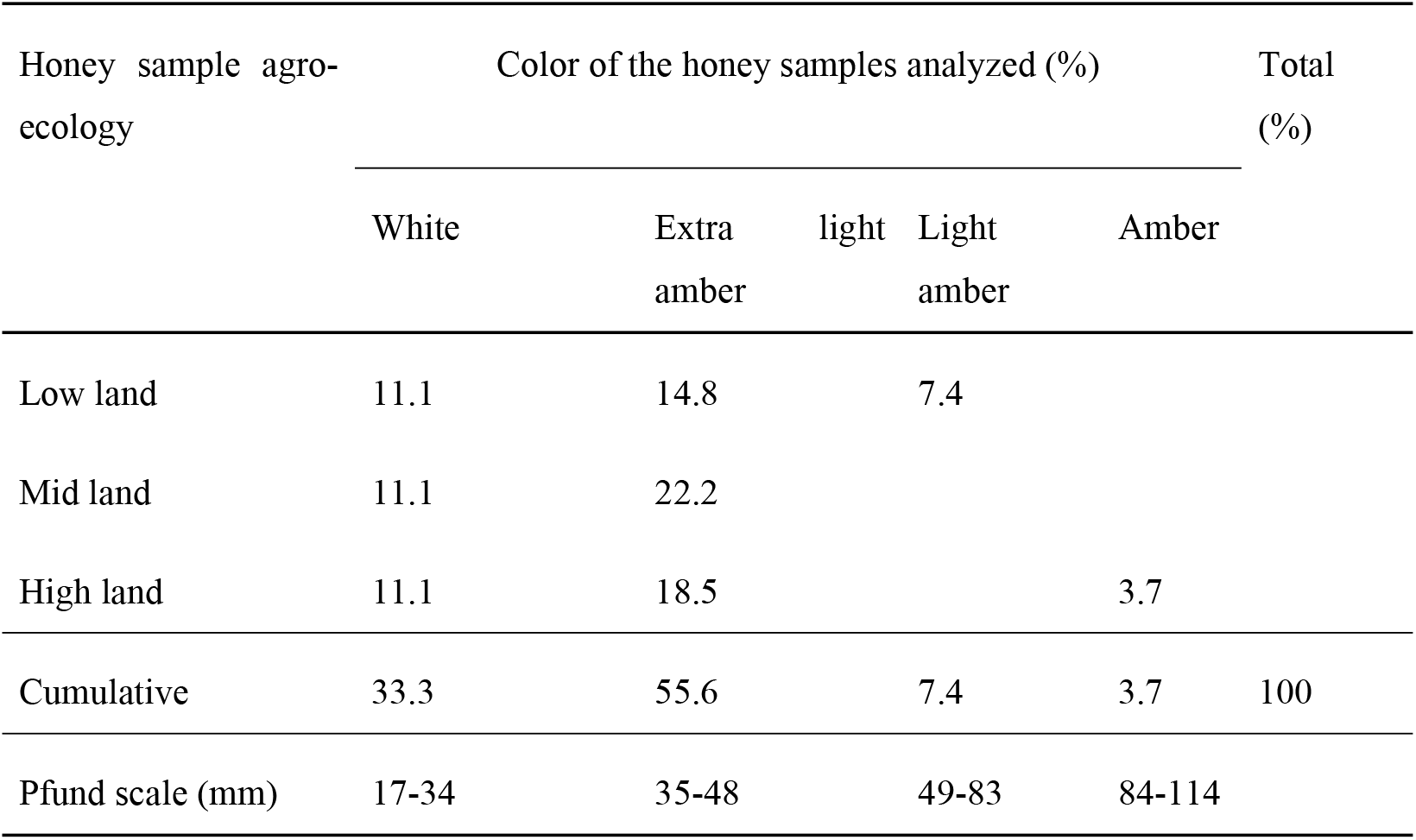
The color of analyzed honey samples obtained from different agro-ecology.

##### B. Moisture content

The honey samples collected from the study area had an average moisture content of 17.16±1.61% ranging from 13.54% to 20.09% (Table 2). There was no significant difference in moisture contents among the honey samples collected from different agro-ecologies and honey collection sources (farmers, SDARC site and honey traders) (p>0.05). Majority of the honey samples collected from the study area (>70.4%) had a moisture content of less than 17.5%.

**Table 2:** Mean physicochemical quality of honey samples obtained from different agro-ecology.

| Agro-ecology | MC (%) | Total solid | pH | FA<br>(meq/kg) | EC<br>(mS/cm) | HMF<br>(mg/kg) | Ash<br>(g/100g) | R.Sugar<br>(g/100g) | Sucrose<br>(g/100g) |
| --- | --- | --- | --- | --- | --- | --- | --- | --- | --- |
| Low land | 16.92±0.58<sup>a</sup> | 83.07±0.58<sup>a</sup> | 3.6±0.21<sup>a</sup> | 21.66±3.56<sup>a</sup> | 0.17±0.05<sup>b</sup> | 2.55±1.99<sup>a</sup> | 0.07±0.03<sup>a</sup> | 67.0±2.26<sup>a</sup> | 4.65±1.10<sup>a</sup> |
| Mid land | 17.34±1.64<sup>a</sup> | 82.65±1.64<sup>a</sup> | 3.58±0.13<sup>a</sup> | 24.21±4.99<sup>a</sup> | 0.27±0.06<sup>a</sup> | 2.46±1.94<sup>a</sup> | 0.12±0.05<sup>a</sup> | 67.02±2.27<sup>a</sup> | 3.62±0.81<sup>a</sup> |
| High land | 17.21±2.3<sup>a</sup> | 82.78±2.3<sup>a</sup> | 3.52±0.13<sup>a</sup> | 26.65±4.79<sup>a</sup> | 0.31±0.09<sup>a</sup> | 0.95±0.79<sup>a</sup> | 0.12±0.05<sup>a</sup> | 65.98±2.79<sup>a</sup> | 4.58±0.87<sup>a</sup> |
| Overall mean | 17.16±1.61 | 82.84±1.61 | 3.56±0.16 | 24.17±4.79 | 0.25±0.09 | 1.99±1.77 | 0.10±0.05 | 66.68±2.41 | 4.29±1.02 |
| P value | 0.863 | 0.863 | 0.529 | 0.082 | 0.001 | 0.092 | 0.062 | 0.588 | 0.051 |
| Minimum | 13.54 | 79.91 | 3.2 | 17.0 | 0.11 | 0 | 0.02 | 61.34 | 2.31 |
| Maximum | 20.09 | 86.46 | 3.87 | 32.97 | 0.43 | 5.53 | 0.24 | 69.92 | 6.18 |
| **CAC(2001)** |  | - | - |  |  |  |  |  |  |
| ***A.mellifera*** | **≤ 20%** |  |  | **≤ 50** | **≤ 0.8** | **≤ 40** | **≤ 0.6** | **≥ 60** | **≤ 5** |
| EU,2002, | $\leq 20\%$ | - | - | $\leq 50$ | | $\leq 40$ | $\leq 0.6$ | $\geq 60$ | $\leq 5$ |
| *A.mellifera* |  |  |  |  |  |  |  |  |  |
| QSAE(2005 | $\leq 21\%$ | - | - | $\leq 50$ | $\leq 0.8$ | $\leq 40$ | $< 0.6$ | $\geq 65$ | $\leq 10$ |
| ) |  |  |  |  |  |  |  |  |  |

Means with different superscript (a, b, c) within the columns are statistically different at $p < 0.05$ .
Note: CAC- Codex Alimentarius Commission, EU- European Union, FA- Free Acidity, EC- Electrical Conductivity, HMF- Hydroxymethyle Furfuraldehyde, MC- Moisture Content, R.Sugar- reducing sugar, QSAE- Quality Standard Authority Of Ethiopia, meq/kg- milliequivalent per kilogram, mS/cm- milliSiemen per centimeter, mg/kg- milligram per kilogram

##### C. pH

The honey samples collected from the study area had an average pH value of 3.56, in which the individual values ranged from 3.2 to 3.87. No significant differences (p>0.05) in pH were observed between honey samples obtained from different sources (farmer, SDARC apiary site, and local traders) (Table 3), and also between honey samples obtained from the different agro ecologies (Table 2).

**Table 3:** Mean physicochemical quality of honey samples obtained from different sources.

| Honey sample source | MC (%) | Total solid | pH | FA<br>(meq/kg) | EC<br>(mS/cm) | HMF<br>(mg/kg) | Ash<br>(g/100g) | R.Sugar<br>(g/100g) | Sucrose<br>(g/100g) |
| --- | --- | --- | --- | --- | --- | --- | --- | --- | --- |
| Farmer<br>(beekeeper) | 17.51±1.6<sup>a</sup> | 82.49±1.6<sup>a</sup> | 3.52±0.11<sup>a</sup> | 25.54±4.27<sup>a</sup> | 0.26±0.08<sup>a</sup> | 0.40±0.6<sup>a</sup> | 0.11±0.04<sup>a</sup> | 67.87±1.08<sup>a</sup> | 4.82±0.99<sup>a</sup> |
| SDARC<br>apiary site | 16.3±1.61<sup>a</sup> | 83.7±1.61<sup>a</sup> | 3.59±0.17<sup>a</sup> | 19.77±1.98<sup>b</sup> | 0.28±0.08<sup>a</sup> | 0.68±1.14<sup>a</sup> | 0.086±0.04<sup>a</sup> | 68.10±1.54<sup>a</sup> | 4.17±0.99<sup>a</sup> |
| Local honey<br>trader | 17.7±1.41<sup>a</sup> | 82.31±1.4<sup>a</sup> | 3.58±0.2<sup>a</sup> | 27.21±4.26<sup>a</sup> | 0.22±0.1<sup>a</sup> | 0.85±1.14<sup>a</sup> | 0.11±0.07<sup>a</sup> | 64.05±1.92<sup>b</sup> | 3.87±0.92<sup>a</sup> |
| Overall mean | 17.16±1.6 | 82.84±1.61 | 3.57±0.16 | 24.18±4.79 | 0.25±0.09 | 0.64±0.97 | 0.10±0.05 | 66.67±2.41 | 4.29±1.02 |
| P value | 0.132 | 0.132 | 0.617 | 0.001 | 0.354 | 0.632 | 0.53 | 0.000 | 0.129 |
| Min | 13.54 | 79.91 | 3.2 | 17.0 | 0.11 | 0 | 0.02 | 61.34 | 2.31 |
| Max | 20.09 | 86.46 | 3.87 | 32.97 | 0.43 | 5.53 | 0.24 | 69.92 | 6.18 |

##### D. Free acidity

The free acidity level of honey samples analyzed in the present study, ranged from 17 to 32.97 meq/kg with a mean value of 24.17meq/kg (Table 2). The free acidity of honey obtained from different agro ecology did not show significant different (p>0.05) but significance difference was observed (p<0.05) between honey samples obtained from different sources, i.e p<0.05 between Site & the Beekeepers (farmers) and between Site & Local traders.

##### E. Ash content

The honey samples in the study area had an average ash percentage of 0.1(g/100g) ranged from to 0.24(g/100g). The results showed no significant difference (p>0.05) between honey samples collected from different agro ecology and sources.

##### F. Electrical conductivity

The overall average electrical conductivity of honey samples analyzed in the current study was 0.25±0.09mS/cm (in which individual values range from 0.11 to 0.43mS/cm). In this study significant difference (p<0.05) was observed in electrical conductivity among different agro ecologies.

##### G. Hydroxyl methyl furfural (HMF)

The mean hydroxymethylfurfural (HMF) content of the study area’s honey samples was 1.99±1.77mg/kg (ranged from 0 to 5.53 mg/kg). No significant difference (p > 0.05) in HMF content was observed between honey samples obtained from different sources (Table 3) and also between honey samples obtained from different locations (agro-ecologies).

##### H. Reducing sugars

The average concentration of reducing sugars (sum of glucose, fructose and maltose) of honey samples analyzed were 67.0±2.26% for lowland, 67.02±2.27% for midland and 65.98±2.79% for highland with mean value of 66.68±2.41% and statistically no significant difference (p>0.05) was observed between all the honey samples collected from different agro-ecologies (Table 2). But significant difference (p<0.05) was observed between the honey samples collected from different sources (farmer, SDARC site and local trader).

##### I. Apparent Sucrose

The average apparent sucrose content of the study area’s honey was 4.29±1.02 % by mass (Table 2). Honey samples collected from different agro ecology exhibited no significant difference (p > 0.05) in their sucrose content (Table 2). The amount of sucrose in the honey samples obtained from the different sources also showed no significant difference (p > 0.05) (Table 2).

### 3.2 Discussion

Honey color (p-Fund mm) The color of honey depends on the origin of the plant, its age, storage conditions, the method of storing, processing, and collecting the honey in old or new combs found in the hive, and the time of day when the honeycomb is formed inside the hive, whether before or after the peak of the honey production (Krell, 1996). Honey color is often reported in millimeters according to the Pfund scale (an optical density reading commonly used in the international honey trade) or the USDA classification (White, 1975c and Crane, 1980). There are several methods for determining the color of honey. The most commonly used methods are based on visual comparison of samples, such as the Pfund, Lovibond, and Jack scales. Honey is usually marketed and graded using the Pfund scale (Bodor and Benedek, 2021).

The color of liquid honey varies from clear and colorless (like water) to dark amber or black. The various colors of honey are basically all yellow and amber. The color varies depending on the plant origin, age, and storage conditions, while the clarity or transparency depends on the amount of suspended particles such as pollen (Eteraf-oskouei T and Najafi, 2012). The most important aspect of honey color is its value in marketing and end-use decisions. Darker honey is more often used for industrial purposes, while lighter honey is sold for direct consumption. In many countries with large honey markets, consumer preferences are determined by the color of the honey (an indicator of preferred taste), so color, along with general definitions of quality, is the single most important factor in determining import and wholesale prices (Krell, 1996).

Moisture is the second most important component of honey. The geographical location where the honey and pollen-producing plants and bee colonies are found, the maturity of the hive, the plant origin of the honey, and the harvesting method (Belay, Haki, ., 2017). These results are consistent with previous reports (Alemu, 2013; Amir, n.d.; Belay, 2017; Abera and Alemu, 2023; Rysha, 2022). The average moisture content of honey in the study area is lower than the national average (20.6%) (Adgaba, 1999).

According to the standards of the Quality Standards Authority of Ethiopia, the moisture content of honey produced in the study area falls within the “A” grade category (QSAE, 2005). According to Ethiopian standards, honey is classified into three categories based on moisture content: Grade A: 17.5–19%, Grade B: 19.1–20%, and Grade C: 20.1–21%. The maximum allowable moisture content of honey reported by the International Honey Commission is 20% (Bogdanov, 2009). Overall, the moisture content of honey collected in the study area was within the acceptable range of national and international standards.

The pH parameter is very important during honey extraction and storage, as it affects the texture, stability and shelf life of honey (Tesfaye ., 2016). The pH of honey can provide a good indicator of the origin of the honey. It can also predict the deterioration of honey during storage (Amir ., 2010). According to French standards (Official Journal of the French Republic, 1977), honey with a pH in the range of 3.5 to 4.5 is obtained from honey. In contrast, Gonnet reported that the pH value of honeydew is between 5 and 5.5, and that of mixtures is between 4.5 and 5.

According to Khalil and (Bogdanova, 2009), flower honey generally has low pH values in the range of (3.3–4.3). In general, the variability in honey pH is due to the harvested plants, bee salivation, enzymatic processes and enzymatic modifications of raw materials (Amir., 2010.). The present study is in agreement with the studies published on Alemu ., 2013), Malaysian honey (Kek ., 2018), and the results were lower than those of Gedebano Gutazer Wolene, honey from central Ethiopia (Gizaw ., 2020), Bale zone of Oromia (Zone ., 2020) and honey from Tigray (Gebremedhin ., 2013). The acidity of honey produced in the study area (24.17 meq/kg) was found to be similar to the results of (Abera and Alemu, 2023; Alemu., 2013) who reported the acidity of honey produced in the United States (24.17 meq/kg) to be 22.3 meq/kg and 23.5 meq/kg, respectively, in Tehulederi and Sekota areas.

The significant differences in free acidity between samples from different sources may be due to differences in storage conditions and potential fermentation. None of the samples exceeded the honey acidity limits recommended by national and international standards. These results indicate the freshness of the honey samples and the absence of unwanted fermentation in the analyzed honey samples.

The average ash content of 0.1 g/100 g was within the acceptable range of national and international quality standards set at 0.6 g/100 g (Codex alimenatarius Commission 2001). The consistency of ash content across sources and agroecologies may be due to the bee species found in the study area and the lack of honey adulteration. *Apis mellifera* Monticolla is the only bee species found in the entire study area. The honey samples obtained from various sources and agroecologies were free of adulteration.

The adulteration of honey with organic substances such as sugar syrup reduces the weight-to-weight ratio of mineral content, as the inorganic compounds in honey are largely replaced by organic adulteration (Abdi ., 2024). Season is also another factor determining the mineral content of honey (Getachew ., 2014). The results of ash content analysis were consistent with those reported by (Alemu ., 2013) which was 0.14%. However, the present value of ash (0.1%) was lower than previous studies which reported 0.22 ± 0.16%, 0.4%, 0.28% and 0.22%, respectively (Getachew ., 2014; Amir ., 2010; Abera & Alemu, 2023;). The mineral content of honey has been found to vary depending on the soil properties, plant species, environmental conditions and origin of flowering plants, which is a direct indicator of climatic and geological pollutants (Berhanu ., 2022).

The average conductivity was 0.25 ± 0.09 mS/cm, which was lower than the value of 0.65 mS/cm reported by (Gizaw ., 2020) and 0.69 ± 0.06 mS/cm (high quality honey collected from natural forests) reported by (Tesfaye ., 2016). However, the present value (0.25 ± 0.09 mS/cm) is in good agreement with the average value of 0.33 mS/cm (range 0.25 mS/cm to 0.39 mS/cm) reported by (Abdi ., 2024).

The electrical conductivity values of the studied honey samples are within the acceptable values of international standards (i.e., less than or equal to 0.8 mS/cm) (Codex alimenatarius Commission, 2001). The significant differences in electrical conductivity between different agroecologies seem to be due to the differences in the botanical origin of the honey. Electrical conductivity has been described as being closely related to the concentration of inorganic salts, organic acids and amino acids. Since the amounts of these compounds found in nectar source plants vary greatly, this parameter is very important to differentiate honey from different floral sources (Gizaw ., 2020; Gela ., 2021).

The low HMF content (1.99 ± 1.77 mg/kg) in honey from the study area is consistent with the value of 1.8 mg/kg (range 0.5–3.2 mg/kg) reported by Development (Alemu ., 2013).), but lower than the results of 48.48 mg/kg reported by (Abera and Alemu, 2023) and 19.52 ± 9.41 mg/kg reported by (Getachew ., 2014).

According to QSAE (2005), the maximum limit for HMF content in honey is 40 mg/kg. The amount of hydroxymethylfurfural in honey is one of the important indicators of honey quality, indicating whether the honey is ripe or overheated (Alemu., 2013). The very low HMF content in the honey samples analyzed in this study indicates that the honey collected in the study area was fresh.

Similarly, Bogdanov (2002) reported that HMF in fresh honey is generally 0 and increases during conditioning and storage depending on pH and storage temperature. Furthermore, Tarasivoulou (1986) reported that the average HMF content of honey produced in Greece increased from an initial value of 0 to 8.8 mg/kg after 1 year of storage.

The average concentration of reducing sugars (66.68 ± 2.41%) was within the acceptable range of national (>65%) and international quality requirements (Codex alimentarius, 2001; EU, 2002), which should be at least 60%. The reducing sugar concentrations obtained in the present study are in agreement with previous results including 66.41%, 66.79 ± 6.96% and 67.3% reported by (Tesfaye., 2016) (Evaluation of Physicochemical Properties of Honey, producer in Bale Natural Forest, South East, 2016); Getachew ., 2014; Alemu ., 2013) is much lower than the result of 74.37% sugar found in Gutazer Wolene, central Ethiopia (Gizaw ., 2020). The significant difference in the reduction of sugar content between different sources suggests differences in the maturity and plant origin of the honey.

Sugar is the main component that determines the properties of honey, and its content is closely related to the maturity and plant origin of honey (Belay; Haki, ., 2017). Honey produced by honeybees is mainly composed of simple sugars, fructose and glucose, with small amounts of sucrose and/or maltose, and trace amounts of non-sugar components (Gizaw et al., 2020).

The average sucrose content of honey from the study area was below the maximum limit of 10% set by QSAE (2005) and 5% set by Bogdanov, 2009, and Codex Alimenatarius Commission, 2001. The average sucrose content of honey from the study area was higher than the national average of 3.6% reported by Adgaba (1999). The sucrose content of honey is used to detect adulteration of honey by adding sugar cane or beet sugar. Consistent sucrose content across different sources and agro ecologies indicates that the honey is natural and pure.

The sucrose content of honey depends mainly on the botanical origin of the honey and, according to international regulatory standards, should not exceed 5% (g/100g), except in some types of honey where this content is naturally higher. Compounds - false acacia (Robinia pseudoaccacia), alfalfa (Medicago sativa), banksia (Banksia menzes), French honeysuckle (Hedysarum), red eucalyptus (Eucaluptus camandulensis), leatherwood (Eucrypis lucida, Eucryphia milliganii), lavender (Lavendula spp.), borage (Borago officinalis) (Codex alimenatarius Commission, 2001).

## 4. CONCLUSION AND RECOMMENDATION

The study on honey from *Apis mellifera monticolla* in the Waghimra zone revealed excellent physicochemical quality, meeting both international (EU and Codex Alimentarius) and national (Ethiopian) standards. Extra light amber honey was the dominant color, representing 55.6% of samples, with all honey samples graded as A. To further elevate Sekota honey as a globally recognized brand, additional research on its quality is recommended. Continuous quality monitoring, along with the development of targeted marketing strategies, could strengthen local market linkages and support potential export opportunities.

